# Arterial pulsations drive oscillatory flow of CSF but not directional pumping

**DOI:** 10.1101/2020.03.13.990655

**Authors:** Ravi Teja Kedarasetti, Patrick J. Drew, Francesco Costanzo

## Abstract

The brain lacks a traditional lymphatic system for metabolite clearance. The existence a “glymphatic system” where metabolites are removed from the brain’s extracellular space by convective exchange between interstitial fluid (ISF) and cerebrospinal fluid (CSF) along the paravascular spaces (PVS) around cerebral blood vessels has been controversial for nearly a decade. While recent work has shown clear evidence of directional flow of CSF in the PVS in anesthetized mice, the driving force for the observed fluid flow remains elusive. The heartbeat-driven peristaltic pulsation of arteries has been proposed as a probable driver of directed CSF flow. In this study, we use rigorous fluid dynamic simulations to provide a physical interpretation for peristaltic pumping of fluids. Our simulations match the experimental results and show that arterial pulsations only drive oscillatory motion of CSF in the PVS. The observed directional CSF flow can be explained by naturally occurring and/or experimenter-generated pressure differences.

## Introduction

The flow of cerebrospinal fluid (CSF) in the brain is hypothesized to play an important role in the clearance of metabolic waste and maintenance of the ionic environment^1–3^. Recent work suggests that the paravascular spaces (PVS) surrounding cerebral arteries provide a low-resistance pathway for the bulk flow of CSF into the brain^1,4–7^. However, this idea that there is bulk fluid movement into the brain is highly controversial, with both simulations^8–10^ and experiments^6,7,11,12^ being put forward both in support of and against bulk flow. One of the leading theories in support of bulk flow in the PVS identifies “peristaltic pumping” as the flow driver, *i.e.,* the idea that heartbeat-driven pulsations pump CSF in the PVS. Peristaltic pumping in a deformable tube is achieved by repeated contractions and dilations propagating along the wall of the tube. In fluid dynamics, peristaltic pumping is a well-understood mechanism of fluid transport. The mechanism of peristaltic pumping of fluids was first demonstrated by Latham^13^. Further work on the peristaltic pumping of fluids has encompassed a wide range of scenarios^14–17^. Calculations made using fluid dynamic principles can make very accurate predictions of fluid flow under peristalsis, and have been used in designing artificial peristaltic pumps^18–20^.

Recent work by Mestre et al.^7^ and Bedussi et al.^6^ used *in vivo* two-photon microscopy^21^ to simultaneously measure arterial pulsations and the flow of CSF in the PVS around the middle cerebral artery (MCA) by tracking the motion of fluorescent microspheres. They found that movement of CSF in the PVS had two components, a constant flow in the direction of blood flow with an average velocity of approximately 20μm/sec, and an oscillatory flow in phase with the arterial pulsations^6,7^, with a peak velocity of approximately 10μm/sec. Based on these observations, it has been proposed that peristaltic motion of the arterial wall generates a “pumping” force that drives the net flow of CSF parallel to the direction of the pulse wave propagation.

In this study, we apply the well-established fluid dynamic principles of peristalsis to study the nature of fluid flow in the PVS, aiming to bridge the gap between experimental observations and hypotheses. As previous studies of CSF flow in the PVS disagree on both the direction and the flow rates^8–10^, we started our calculations by revisiting the mechanism of peristaltic pumping using time-dependent fluid dynamic simulations with fluid particle tracking in a deforming domain. By emphasizing the mechanism of peristatic pumping, we aimed at providing a clear physical interpretation for our calculations. We then performed fluid dynamic simulations on more realistic models of the PVS. Our simulations suggest that peristalsis with physiologically-plausible pulsation cannot drive the experimentally-observed fluid flow. However, we found that a small, constant pressure gradient (of order 0.01 mmHg/mm) can account for the net forward movement observed experimentally. These results suggest that the observed directional movement of CSF in the PVS is generated by naturally occurring and/or experimenter-generated pressure differences, but not by arterial pulsations.

## Results

We first examine how peristaltic motion affects the flow of an incompressible fluid in a two-dimensional tube. Consider a fluid-filled tube with deformable walls and no pressure difference across its two ends. When the position of the walls is fixed, there is no pressure gradient, and therefore no fluid flow (Fig 1a, 1b). When the walls move inward due to a peristaltic wave propagating to the right (Fig 1a), the fluid-filled domain deforms and the fluid is displaced. When the direction of the fluid flow is the same as the peristaltic wave, the motion is said to be anterograde – otherwise it is said to be retrograde. The flow in both directions is a result of the fluid pressure distribution, shown in Figure 1b. The pressure is maximum at the location of the moving neck and is minimum at the two ends of the tube. Therefore, the fluid that is displaced by the wall is subject to the same pressure difference (Δ*p*) in either direction (Δ*p_a_* = Δ*p_r_*, where the subscripts ‘a’ and ‘r’ denote the anterograde and retrograde flows, respectively). However, since the width of the tube (h) is smaller in one direction (*h_a_* > *h_r_*), there is more resistance for retrograde flow than anterograde flow (since flow resistance R scales with width of the tube *R* ∝ 1/*h*^3^ for 2D flow, *R_a_* < *R_r_*). As a result of this difference in flow resistance, the anterograde flow is greater than the retrograde flow (flow rate, *Q* = Δ*p/R, Q_a_* > *Q_r_*). Thus, while peristalsis drives both anterograde and retrograde flows, a net flow in the direction of the peristaltic wave emerges (Video SV1). This example is a simplified version of peristalsis, where, the walls of the fluid-filled tube only contract. An example with a periodic contraction and expansion of the walls is demonstrated in figure S1 and Video SV2.

**Fig 1.**
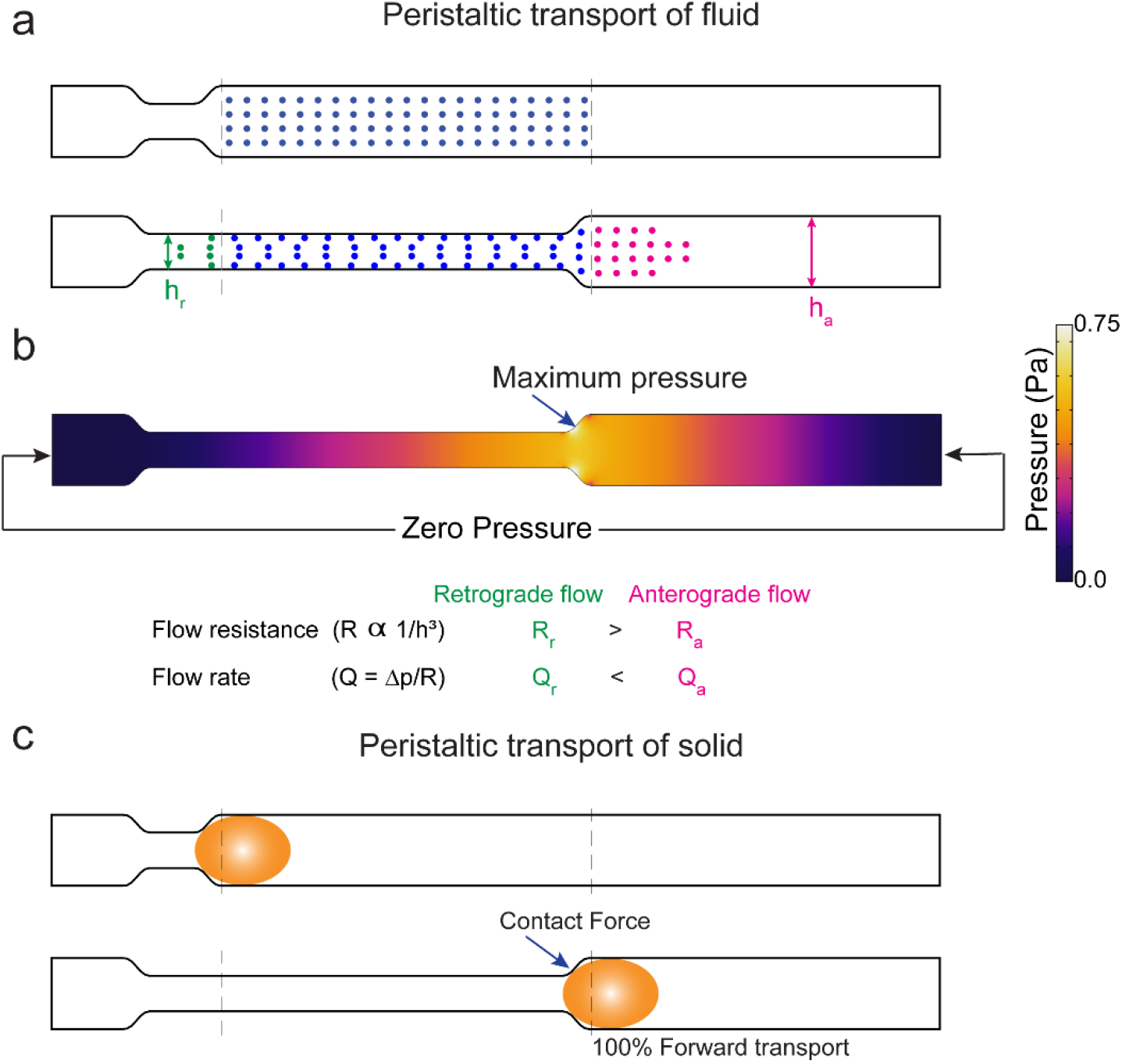
Mechanism of forward transport (pumping) driven by peristalsis in fluids and solids. **a.** The movement of fluid (the dots represent fluid particles) driven by peristaltic compression of the walls of a 2D tube. The fluid movement in this figure is calculated using the Navier-Stokes equation with zero traction (pressure) at the two ends of the tube. **b.** The pressure field in the tube in the deformed state. The fluid is displaced by the moving walls and this creates high pressure at the neck of wall movement. The pressure difference is same for retrograde and anterograde flow. However, the resistance is large posterior to the site of contraction. Compared to anterograde flow, retrograde flow needs the fluid to flow through a narrower tube. This results in a greater anterograde flow (magenta dots in **a**) compared to retrograde flow (green dots in **a**). **c.** The movement of a solid bolus (yellow ellipse) driven by peristaltic motion of the walls of a 2D tube. The contact forces between the moving walls and the solid bolus are responsible for forward transport. The position of the solid bolus with respect to the dotted lines shows that the solid is moved forward.

It is important to bear in mind the difference between peristaltic transport of fluids and peristatic transport of solids. The textbook picture of peristalsis^22,23^ is derived from the transport of solid matter in the esophagus and the gastro-intestinal tract. When solid matter is transported by peristalsis, all of the material moves in the direction of the peristaltic wave (Fig 1c). This differs from the case of fluid transport by peristalsis, which generates both anterograde and retrograde flows (Fig 1b, S1). Moreover, the peristaltic transport of solids is independent of the magnitude of wall motion and the length of the tube. In contrast, the nature of fluid flow in a tube driven by peristaltic motion of the walls is highly depended on the magnitude of both wall motion and tube length^24^, which we will demonstrate in the results. This understanding of the mechanism of peristaltic transport of fluids is crucial to interpreting the results of fluid dynamic models of the PVS. The assumptions of the shape, size and the deformation of the PVS may vary between the models, but the mechanism of peristaltic transport remains the same.

### Peristaltic pumping requires unphysiologically large amplitude pulsations for meaningful fluid flows

To understand the relation between arterial wall movement and fluid movement in the PVS, we created a model of peristaltic pumping. In our model, the geometry of the PVS is taken to be cylindrically symmetric, with the artery centered within the PVS (Fig 2a). While the geometry of the fluid domain is simplified, the inner and outer radii are based on realistic values (see methods). We then imposed a sinusoidal peristaltic wave on the arterial wall, while keeping the outer wall of the PVS fixed, effectively making the brain tissue rigid. In order to capture the whole peristaltic wave, the length of the PVS used in the simulation was equal to one wavelength (λ) of the peristaltic wave (see methods). Since we are interested in studying the pumping generated by arterial wall movement alone, we used periodic boundary conditions at the axial ends of the PVS. This is equivalent to studying flow driven by peristalsis with no additional pressure differences^8,14,15,25^. We tracked the motion of particles at the center of the PVS.

**Fig 2.**
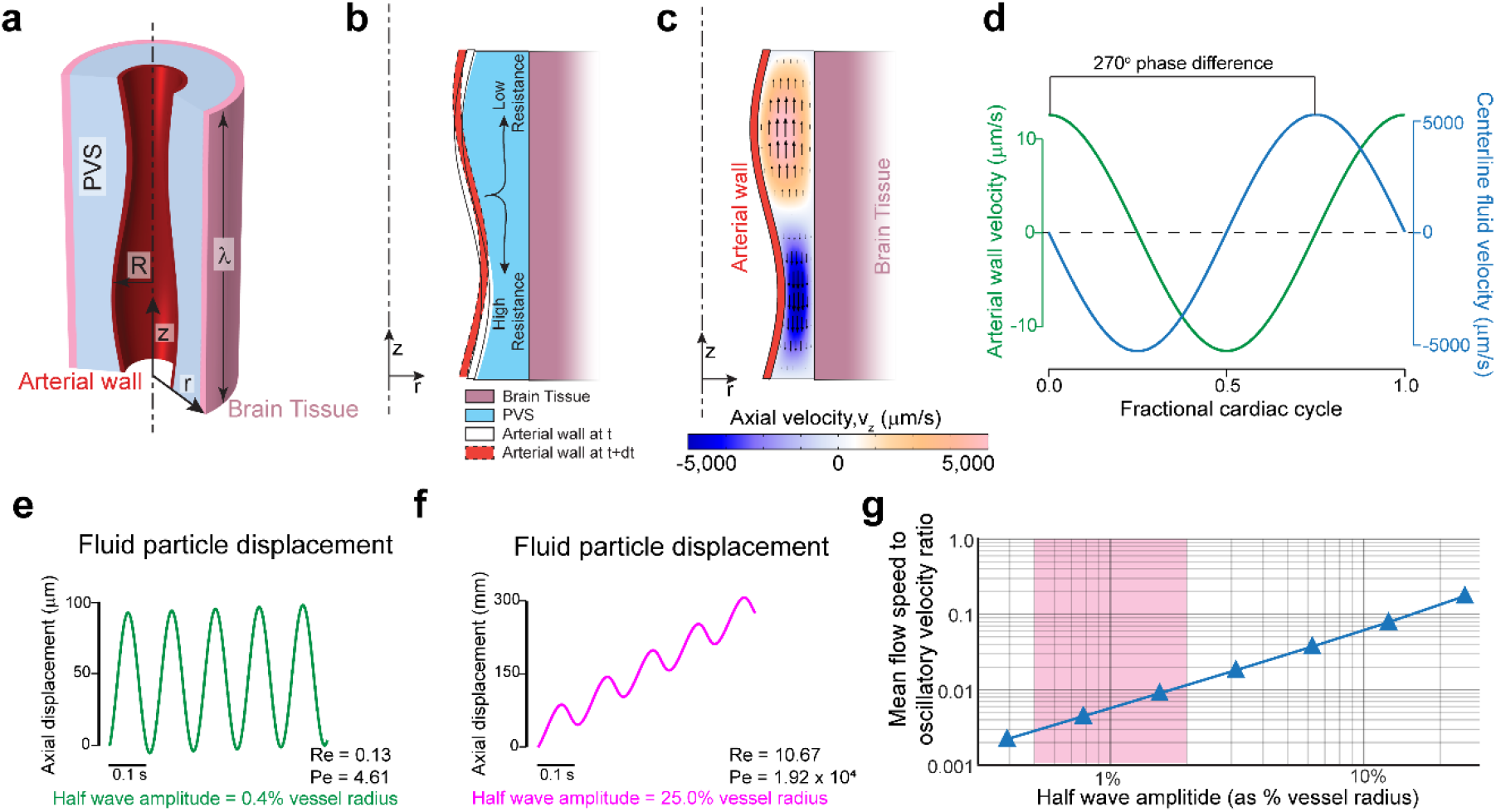
Heartrate pulsations drive oscillatory’ but not directional flow. Large non-physiological pulsations are required for appreciable peristaltic pumping. **a.** Schematic of the axisymmetric peristatic pumping model. The arterial wall undergoes peristatic movement, while the outer wall of the PVS is fixed. The length of the PVS is taken as one wavelength of the peristaltic wave (λ) with periodic boundary conditions at the two axial ends. **b.** Schematic of the fluid motion during periodic peristalsis. The fluid displaced by the moving wall should move with lower resistance along the direction of the peristaltic wave than in the opposite direction. This difference in flow resistance results in net forward pumping. **c.** The magnitude of the axial velocity is denoted by color, with arrows showing the direction. The half-wave amplitude of the peristaltic wave is 0.8% of the vessel radius. The deformations are scaled by a factor of 50 in the figure to clearly show the arterial wall movement. **d.** The plots show the relative phase between arterial wall velocity and the centerline fluid velocity at the same axial (z) location (midpoint, z = λ/2). There is a 270° phase difference between the wall velocity and fluid velocity. **e.** The trajectory (in z) of a fluid particle at the center of the PVS, where the amplitude of the peristaltic wave is similar to heartbeat driven pulsation^6,7^ (0.8% peak-to-peak change in arterial radius). The flow is mostly oscillatory with very little unidirectional movement. **f**. The trajectory (in z) of a fluid particle at the center of the PVS, where the amplitude of the peristaltic wave is unrealistically large (50% peak-to-peak change in arterial radius). This trajectory with appreciable unidirectional movement looks similar to experimental results^6,7^. **g.** Plot showing the ratio of mean flow speed (average unidirectional flow velocity) to oscillatory velocity (peak velocity change in a cycle) as a function of the amplitude of the peristaltic wave. The shaded region shows the normal amplitude of pulsation cerebral arteries (1-4% of arterial radius peak-to-peak). Note that in this range, the oscillatory velocity is ~ 2 orders of magnitude higher than the pumping velocity.

Peristaltic pumping of fluid is a result of lower flow resistance for anterograde flow and higher resistance for retrograde flow (see Fig 1 and Fig 2b). This explains the fluid velocities observed with respect to the arterial wall position and wall velocity (Fig 2c, 2d). The phase difference between the arterial wall velocity and the downstream fluid velocity (axial velocity, *v_z_*) remained the same throughout the length of the domain. Flow resistance (*R_flow_*) of a tube with an annular cross-section decreases with approximately the fourth power of the internal radius^26^.

For slow, laminar flows like those in the paravascular space, the flow resistance of a tube with annular cross-section is given by the equation:

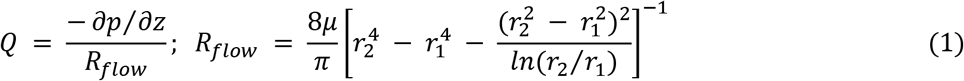

Where Q is the flow rate, p is the pressure and μ is the dynamic viscosity of the fluid. The internal and external radii of the annular region are given by r_1_ and r_2_ respectively. For our calculations, Reynolds numbers range from 0.13 to 10.67, well within the laminar flow regime. Given the strong dependence of fluid resistance on the diameter, it follows that the amplitude of pulsations (the change in internal radius) should have a large effect on the flow resistance changes and therefore the pumping generated by peristaltic motion.

We examined the relation between pulsation amplitude and the trajectory of the fluid particles in the PVS. To put our results in the context of experimental findings, the typical half wave amplitude of heartbeat pulsations is 0.5-2% of arterial diameter^6,7^. Our simulations show that such small amplitude pulsations generate little difference between forward and backward flow resistance, which resulted in oscillatory fluid flow with minimal net anterograde flow (Fig 2e). These simulations show that the mean downstream velocity for fluid in the PVS driven by heartbeat pulsations should be approximately 2 orders of magnitude smaller than the oscillatory velocity. However, the kind of fluid particle trajectories reported by Bedussi *et al*.^6^ and Mestre *et al.*^7^ are very different from the ones simulated in Fig 2e. The fluid trajectories in the PVS observed in both studies are more similar to the ones shown in Fig 2f, where the net anterograde motion of the fluid is of the same order as the oscillatory motion. This kind of fluid motion would require non-physiological amplitudes arterial pulsations, with half wave amplitudes around 25% of the arterial radius. To better understand the effect of pulsation amplitude on fluid flow, we examined the relation between the pulsation amplitude and the ratio of mean flow speed (or average anterograde velocity) to oscillatory velocity (difference between peak anterograde velocity to peak retrograde velocity) (Fig 2g). These result show that heartbeat-driven pulsations in an idealized model are too small to explain the directed flow of CSF seen *in vivo.*

While the shape of the PVS in our model was simplified, the model still provides important generalizable insights into the mechanism of peristalsis. Specifically, the model helps us understand the relation between the movement of the arterial wall and the flow of fluid. We found that the radial wall velocity and the anterograde fluid velocity are always out of phase (by 270°, Fig 2c, 2d), and that the kind of fluid trajectories observed *in vivo* would require large, non-physiological amplitudes for arterial pulsations. Next, we examined if these results held for a 3-dimensional model of peristalsis with a realistic shape of the PVS and pulse waveform.

### Paravascular flow measurements are inconsistent with peristaltic pumping

To test if fluid flow is influenced by the details of the shape of the PVS or the waveform of heartbeat driven pulsations, we created a model with a realistically-sized and shaped PVS, with a cardiac waveform drawn from experimental data^7^ (Fig 3a,b). The outer wall of the PVS was assumed to be fixed, and the length of the domain was set to be equal to one wavelength (*λ*) of the peristaltic wave. We use a no pressure (traction) boundary condition at the boundaries in place of the periodic boundary condition used for the axisymmetric simulations. This is done to better estimate the pressure changes in the PVS (Fig 3e).

**Fig 3.**
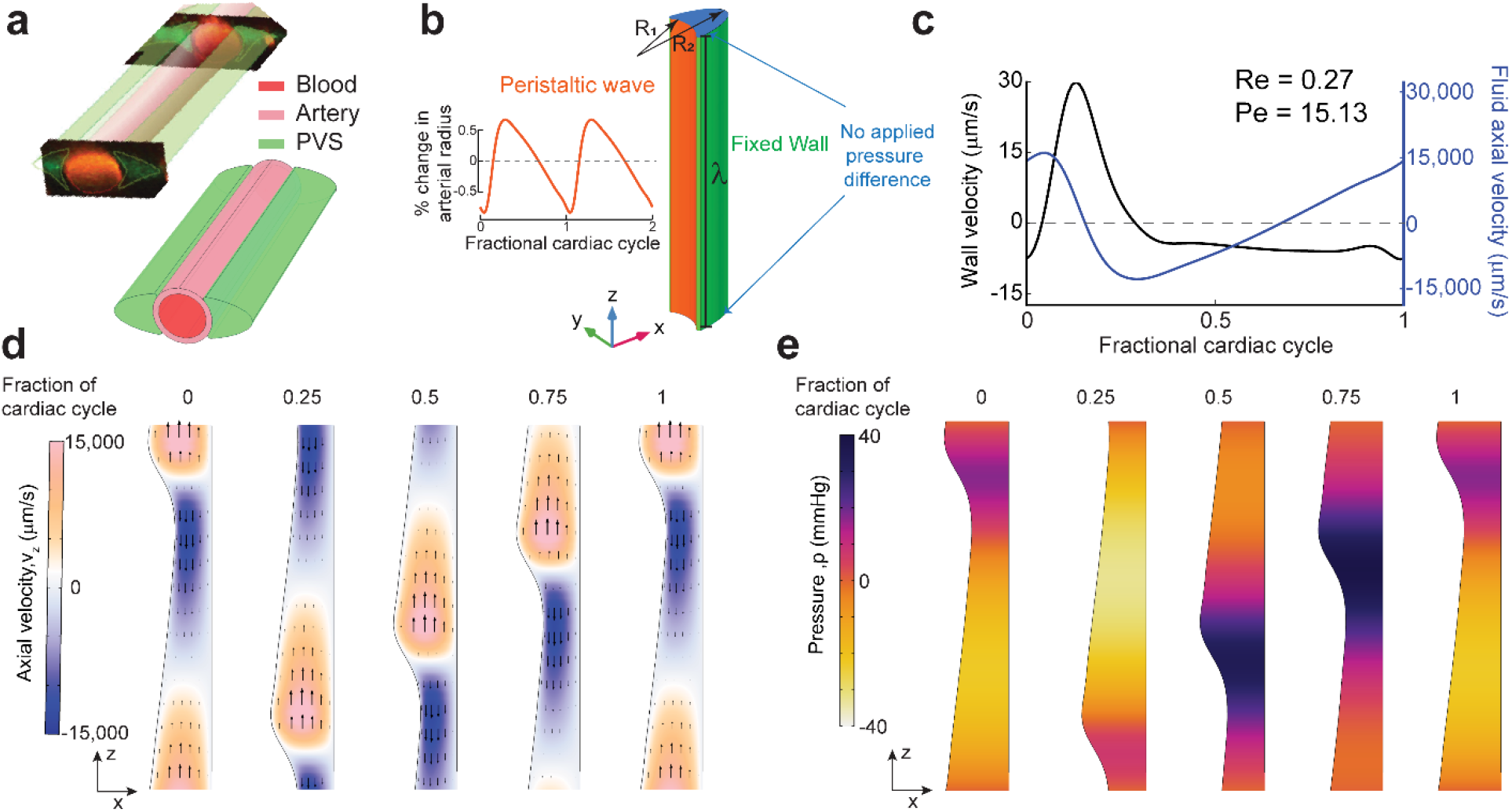
Simulations of arterial pulsations in a model with an elliptic PVS show that observed fluid flows are inconsistent with a peristaltic pumping mechanism. **a.** The 3D geometry of the PVS used in our simulations replicates the PVS geometry observed *in vivo* in mice. The figure in top-left is taken from Mestre *et al.*^7^ and the figure on the bottom-right shows the 3D geometry used in our calculations. **b.** The dimensions and boundary conditions used in 3D fluid dynamics simulations. The wall movement on the arterial side (orange) is given by a travelling wave with a realistic waveform observed *in vivo* in mice^7^ (inset). The wall on the brain tissue side is fixed (green). The parameters (R_1_, R_2_, λ etc are given in Table 1). **c.** Plots showing the arterial wall velocity and the centerline velocity of the fluid taken at the same axial (z) location. While the wall velocity profile and magnitude are very similar to what was observed *in vivo*^7^, the oscillatory fluid velocity is ~1000x higher than the values observed *in vivo*^6,7^. Moreover, the peak fluid velocity is not in phase with the wall velocity. These discrepancies were predicted by our simplified model (Figure 2). **d.** The colors show axial velocity profile at the mid-section of the PVS (XZ plane at y=0) throughout the cardiac cycle. Arrows are provided to make the interpretation of flow easier. The fluid velocity profile agrees with our expectations from the mechanism of peristalsis (Fig 2b) and our simplified model (Fig 2c). The deformations are increased by a factor of 50 in post-processing to clearly show the arterial wall movement. **e.** Corresponding pressure profile at the mid-section of the PVS (XZ plane at y=0) throughout the cardiac cycle. The deformations are scaled by a factor of 50 in post-processing to clearly show the arterial wall movement.

The mean flow speed (anterograde velocity time-averaged over a complete cycle) of fluid particles at the centerline of the PVS in our simulation was 102.1 μm/s. However, this was accompanied by oscillatory fluid velocities of approximately 30,000 μm/sec well over a hundred times the mean flow speed (Fig 3c, 3d). Moreover, in our simulations the fluid downstream velocity and the arterial wall velocity were out of phase, whereas these velocities were in phase for *in vivo* measurements by both Bedussi *et al*.^6^ and Mestre *et al*.^7^ (Fig 4d). Viewed together with the results of the axisymmetric simulations, our simulations suggest that the shape of the PVS and the waveform of heartbeat pulsations cannot pump CSF in a model with a simplified geometry of the PVS.

**Fig 4.**
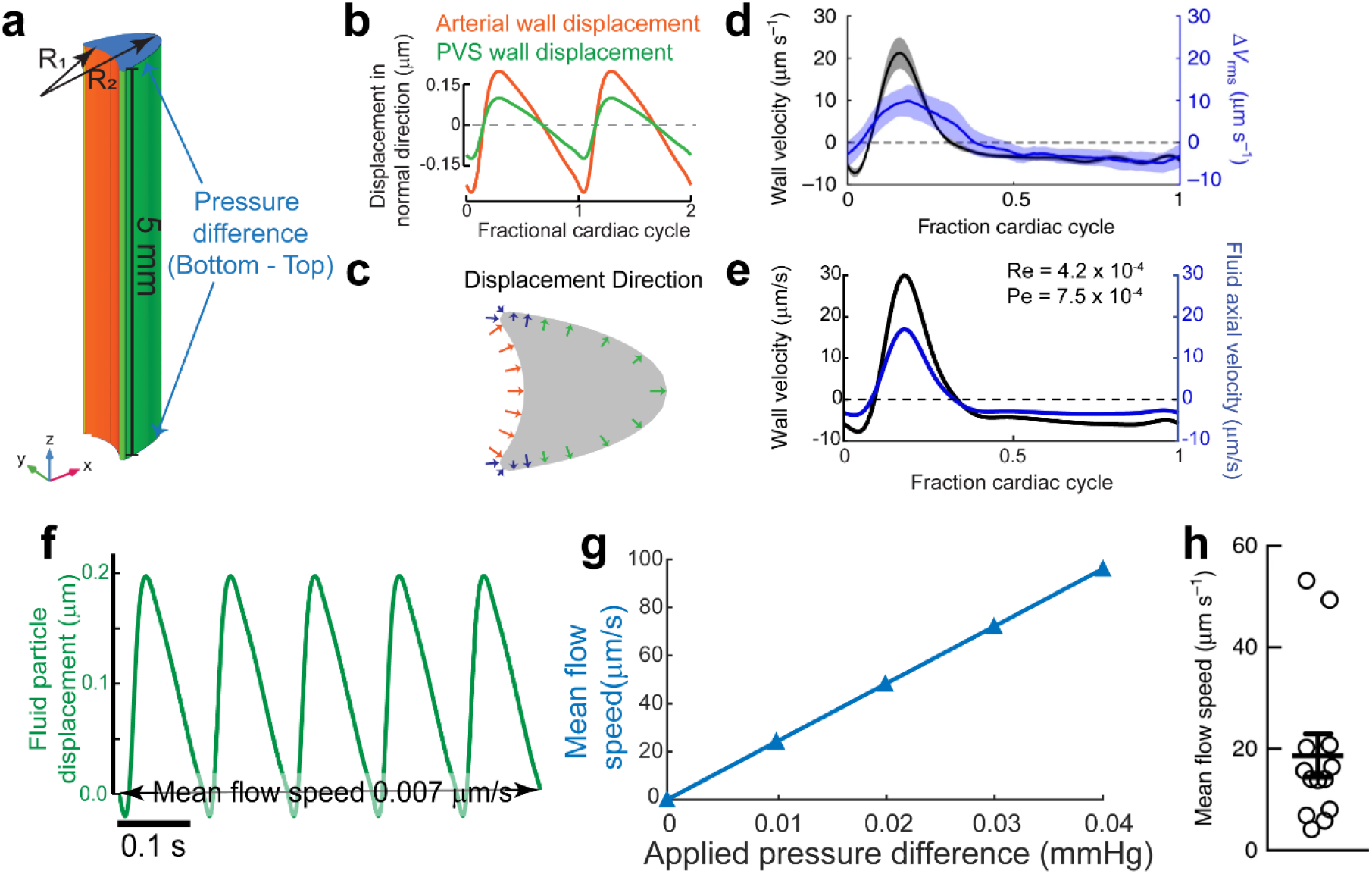
A small, constant pressure gradient can explain directed fluid movement *in vivo*. **a.** Schematic of the model. The length of the PVS is set to 5 mm to match the length of the MCA in mice (4-6mm^6,32,33^). An additional pressure difference between the inlet and outlet is applied. **b.** The displacement of the arterial wall (orange) and the PVS wall (green) used in the simulation. The displacement is given in the direction of the surface normals shown in c. **c.** Positive displacement direction at a cross-section of the PVS, this is the direction of displacement for the plots shown in **b**. The Inner wall is shown in orange and the outer wall is shown in green. The displacement changes in amplitude and direction from the inner wall to the outer wall. This transition is carried out using the smooth, step function in COMSOL Multiphysics. The changing direction and length of the blue arrows indicates the smooth transition. **d.** Plot showing arterial wall velocity and oscillations in fluid velocity measured *in vivo.* Adapted from Mestre *et al.* ^7^ **e.** Plot showing arterial wall velocity and oscillations in downstream fluid velocity our simulation matches with those in **d**. **f.** Plot of the trajectory of a fluid particle in the z direction shows that there is very little pumping by arterial wall movement. **g.** Plot showing the relation between an applied pressure difference across the ends of the PVS and the mean anterograde flow speed. An additional 0.01 mmHg pressure difference across the PVS is required to achieve a mean flow speed of 24.4 μm/s, similar to the mean flow speeds observed *in vivo* (**h**). **h.** Mean flow speed in the PVS measured *in vivo.* Adapted from Mestre *et al.*^7^

These results show that a peristaltic pumping model is inconsistent with experimental findings, which suggests that there are some problems with the assumptions of the peristaltic pumping model. In the next section, we revisit these assumptions and attempt to match the results of the fluid dynamic calculations with experimental findings.

### Pressure differences, not arterial pulsations drive bulk fluid flow

In order to better capture the geometry of the PVS, we made changes to our 3D model based on the anatomy of the brain, the subarachnoid space and cerebral vasculature. We shortened the length of the PVS to 5 mm, and made the outer wall of the PVS move with the pulsations. Previous peristaltic pumping models^8,9,14,15^ have set the length of the fluid chamber to be equal to one wavelength of the peristaltic wave. However, the wavelength of the peristaltic motion of arteries is considerably larger than the length of the middle cerebral artery (MCA), the proposed source of peristaltic pumping. With a peristaltic wave speed of 0.5-2 m/s^27,28^, and a heartbeat frequency of 6-10 beats/second in mice^6,7,29–31^, the wavelength of the peristaltic wave is between 50-160mm in mice, while the MCA is only 4-6 mm long in mice^6,32,33^. This means that the pulse wave travels so fast across the length of the PVS that the arterial wall moves in and out simultaneously and therefore there is no appreciable difference in the flow resistance for anterograde and retrograde flows (Fig S2). We calculated the fluid particle trajectories 1 mm from the distal end of the PVS segment (z = 4 mm), which captures the geometry of the surface of the brain where the flow measurements were made. This corrected the inconsistency between the phase of the fluid downstream velocity and the arterial wall velocity found in the peristaltic pumping model with a length of one pulsation wavelength (Figure S2). Our results are similar to the phase relation between arterial wall velocity and fluid velocity estimated by Asgari *et al*._10_, who studied the flow driven by penetrating arterioles in a model with anatomically realistic dimensions.

Secondly, the fixing of the outer wall of the PVS in other models means that the brain tissue and the subarachnoid space that surround the PVS are rigid. This is not realistic because the brain tissue is very soft, with a shear modulus in the range of 1-8 kPa^34–38^ (7.5-60 mmHg) and the 80 mmHg pressure changes (Fig 3e) predicted by the peristaltic pumping model will cause substantial deformations. To include the effect of the soft tissue, we moved outer wall of the PVS with the same frequency as the heartbeat driven pulsations (Fig 4b). We applied these small, pressure-driven deformations in the direction of the outward normal of the surface of the PVS, because pressure-like forces act along the outward normal of a surface. Since the mechanical properties of the subarachnoid space are mostly unknown^39–42^, we adjusted the amplitude of these deformations so that the oscillatory fluid velocity matched that observed *in vivo* by Mestre *et al*.^7^

Our simulations suggest that the wall movement itself can only drive oscillations in fluid flow with negligible (0.007 μm/s) mean anterograde flow. The time course of fluid velocity from our simulations and its relation to the arterial wall movement agrees very well with the measured values in both phase and magnitude (Fig 4d and 4e). The phase relation between the arterial wall velocity and the fluid velocity is a direct result of correcting the length of the PVS, while the magnitude of fluid velocity is corrected by including movement of the outer walls of the PVS in the simulation. However, the simulations suggest that arterial pulsations generate very little net anterograde flow with a time averaged downstream velocity of 0.007 μm/s (Fig 4f).

Finally, we tested the possibility that small pressure differences across the ends of the PVS can drive the bulk flow observed in the experimental studies. We calculated the fluid flow through the PVS while varying the imposed pressure difference over a physiologically plausible range. We found that a very small pressure difference, 0.01 mmHg across the length of the PVS (5 mm), was sufficient to drive a mean downstream speed of 24.4μm/sec (Fig 4g), close to the mean flow speed observed *in vivo*^7^(Fig 4h). The pressure difference value was also found using equations derived in another recent theoretical study that estimated the flow resistance of paravascular spaces^26^. Such a small pressure difference is practically impossible to measure in live animals, due to the lack of instruments sensitive to such small changes^43,44^. The pressure differences could be normally present due to CSF production in the ventricles^43^ and drainage via meningeal lymphatic vessels^45,46^ and the cribriform plate^47^, or be generated by intracranial injections^48,49^ of the tracer spheres. We conclude that peristalsis cannot drive unidirectional fluid pumping in the PVS of cerebral arteries under physiological conditions and that the experimentally observed CSF flow in the PVS is probably due to pressure differences present in the system.

## Discussion

Peristatic pumping has been hypothesized to drive directed movement of cerebrospinal fluid in the paravascular space. In this study we test the “peristaltic pumping” hypothesis, by using simulations of fluid dynamics to understand what experimental measurements tell us about bulk flow. We started with simple models to provide a physical interpretation to the process of peristalsis of fluids and built more physiologically realistic models informed by the results of these models. We were able to improve upon previously published computational models aimed at studying the flow of CSF in the PVS^8–10,25^, using the detailed anatomical and physiological information from the experiments by Mestre *et. al.*^7^. This experimental data provided information about the shape of the PVS around cerebral arteries and the amplitude and waveform of the heartbeat driven pulsations, which we used in our modeling. The experiments also had detailed information on the oscillatory and anterograde flow of CSF in the PVS. Our simulations show that the cardiac pulsation of arteries is only capable of driving the oscillatory motion of CSF in the PVS, and not the unidirectional bulk flow. Rather, the experimentally observed unidirectional flow is likely to be driven by pressure differences in the system.

Our simulations point to two main reasons why arterial pulsations cannot drive unidirectional fluid flow in the PVS. First, direct measurement of cortical arteriole diameters in mice using two-photon imaging shows that the amplitude of the heartbeat-driven pulsations is small (1-4% peak to peak change in arterial diameter^6,50,51^). In humans, CT angiography has shown that pulsations drive only a maximum of 4-6%^52,53^ change in the volume of the MCA (2-3% change in diameter assuming a cylindrical geometry). Our calculations show that substantially larger cardiac pulsations (roughly 50% peak-to-peak change in diameter) are required to drive significant directed motion of the fluid relative to the oscillatory motion. Second, the peristaltic motion of arteries cannot drive unidirectional fluid flow because the length of the PVS is substantially less than the wavelength of the peristaltic wave. The total length of the MCA is between 4-6 mm in mice^6,32,33^, while the wavelength of the peristaltic wave is between 100-1000 mm (based on the pulse wave velocity of 1-5 m/s^27,28^ and a heart rate of 6-12 Hz^29,30^). This is over an order of magnitude difference between the length of the PVS and the wavelength of the peristaltic wave. In humans, the MCA is longer, (roughly 100 mm^54^). However, while the pulse wave velocity, a function of arterial stiffness^55,56^, remains roughly the same in mice and humans^57,58^, the heartrate in humans is around 1-2 Hz, which makes the wavelength of the peristaltic wave 1-2 orders of magnitude higher than the length of the MCA in humans. Therefore, in mice as well as in humans, arterial pulsations are unlikely to drive unidirectional CSF flow.

Based on the experimental evidence available, we speculate that two possible mechanisms that could drive CSF flow in the PVS, namely, CSF production in the choroid plexus and osmotic pressure differences across astrocytic end feet. CSF flow through the PVS and into the brain is severely affected in aquaporin-4 (AQP4) knockout mice^1,50^. The AQP4 channel is selectively permeable to water^61,62^ and is present in the choroid plexus^63^ and the astrocytic endfeet^1^. The deletion of the AQP4 gene could reduce CSF production and osmotic flow through astrocytic endfeet. It is possible that a combination of the two factors drive CSF flow since the osmotic concentration gradients and the CSF production rate are interrelated^69^. Alternatively, the observed flow in the PVS might be an caused by the infusion rate of 1-2μl/min used in the experiments to study CSF flow^6,7^ which is 3-5 times the typical rate of CSF production rate in mice (0.38 μl/min^43^). The infusion rate used in these experiments is known to increase intercranial pressure^1,44^, as pointed out by Hladky and Barrand^48^. A detailed 3-D model of the whole brain with the PVS and the SAS, all modelled as poroelastic media^70^ would be needed to test the possibility of the observed flow being an artifact of the infusion.

An important result of our simulations is that the paravascular spaces around pial arterioles provide a crucial pathway for fluid transport in the brain due to their low flow resistance. A very small pressure difference (0.01 mmHg, Fig 4g) across the length of the MCA (5 mm) can be sufficient to drive fluid through the PVS with a mean speed of ~20 μm/s. This is in stark contrast to the much less permeable brain tissue, where a pressure gradient of 1 mmHg/mm can only generate fluid velocities in the order of 0.010 μm/s^59^. However, the low flow resistance makes understanding the driving force for CSF movement in the PVS extremely difficult. A pressure difference in the range of 0.01 mmHg cannot be accurately measured with current instruments, which have a resolution of around 1 mmHg^43,60^. Moreover, invasive access of the skull probes through the skull severely affects the flow through the PVS^50^.

## Methods

### Model equations and boundary conditions

We use a standard time-dependent finite element method to solve the equations of fluid motion in the PVS. These equations are formulated to correctly account for the deformation of the PVS. Specifically, we write the equations in Arbitrary Lagrangian-Eulerian (ALE) coordinates (see appendix). As is well-known, ALE formulations are able to account for the deformation of the solution’s domain at the expense of having to determine an auxiliary motion typically referred to as the “mesh motion”^71–73^. The governing equations for the fluid and the mesh movement are written in their weak, tensor form (see Appendix) and converted to their component form using Wolfram Mathematica. These component form equations are implemented in COMSOL Multiphysics (Burlington, MA) using the “Weak Form PDE” interface, where PDE stands for partial differential equation. Therefore, the overall solution scheme is our own, and COMSOL Multiphysics simply provides a high-level integrated programming environment within which said scheme is implemented.

The fluid (CSF) velocity and pressure are governed by the incompressible Navier-Stokes equations (eq. M1 – M3). We solve for the fluid velocity (***ν***_*f*_) and pressure (*p_f_*) in the PVS as a function of time (*t*). In eq. M1, *ρ_f_* and ***σ**_f_* are the fluid’s mass density and Cauchy stress, respectively. In eq. M3, *μ_f_* is the fluid’s dynamic viscosity.

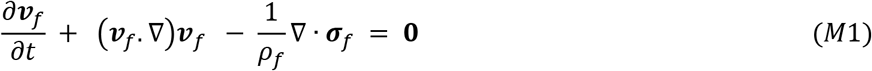

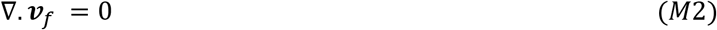

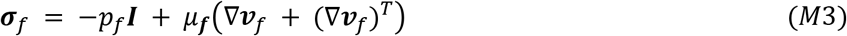

The governing equation for the mesh motion is dictated by convenience and, where necessary, by the problem’s geometric constraints. In our problem, the deformation of the solution’s domain (PVS) is relatively mild and therefore we the mesh motion equation, with primary unknown given by the mesh displacement ***u**_m_*, is chosen to be a linear elliptic model^74^, namely the Laplace equation (eq. M4):

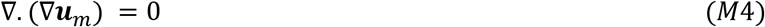

We use no-slip boundary condition at the inner and outer walls of the PVS, *i.e.*, fluid velocity is equal to the wall velocity in all simulations (eq. M5). For the axisymmetric simulations, the inner walls have a baseline radius of R_1_ and the outer walls have a fixed radius of R_2_. The movement of the inner walls is given by a travelling sinusoidal wave (eq. M6). There is no wall movement at the outer wall (eq. M7). The total length of the tube is taken equal to the wavelength (*Λ*) of the peristaltic wave. Periodic boundary conditions are used at the two ends of the tube (eq. M8). To obtain a unique pressure solution, a global constrain is applied for the total pressure (eq. M9).

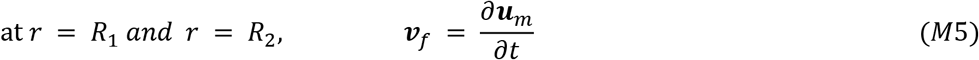

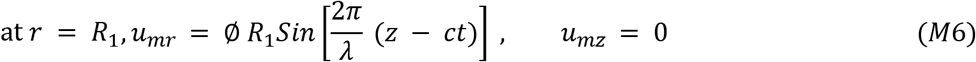

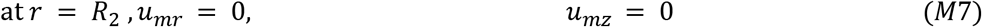

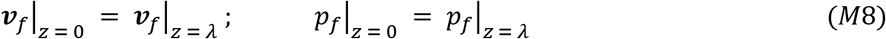

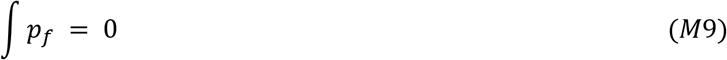

In eq. M6–M7, *u_mr_* and *u_mz_* are the r and z components of the mesh displacement (***u**_m_*). ø is the half wave amplitude of the peristaltic wave, as a fraction of the baseline diameter R_1_. c is the speed of the peristaltic wave. The integration in equation M9 is performed over the entire computational domain.

For the 3D simulations presented in Fig 3 and 4, we created the cross section of the PVS to resemble the geometries observed *in vivo*^7,26^. The inner wall of the cross section is a circle of radius R_1_. The outer wall of the cross section is an ellipse with major axis R_2_ and minor axis 0.8R_1_. The intersection of the circle with the ellipse is smoothened with a fillet of radius 0.08R1. The cross-section can be divided into three regions. The three regions can be identified in Fig 4c. The inner walls of the PVS (the walls facing the arteries or the circular face) are shown in orange. At the inner walls, a dilation of the arterial wall will cause a deformation of the PVS in the direction opposite to the unit outward normal, ***n*** (eq. M10). The outer walls of the PVS (wall facing the SAS or the brain tissue or the elliptical face) are shown with green arrows in Fig 4c. On the outer walls, the pressure is higher when the vessel dilates and lower when the vessel contracts (Fig 3e). Therefore, when the vessel dilates, the ougter walls of the PVS deform in the direction of the outward normal, ***n*** (eq. M11). We call the part of the wall between these two regions, the transition region (the fillet region shown with blue arrows in Fig 4c). Here, the displacement is smoothly transitioned using the step function available in COMSOL Multiphysics.

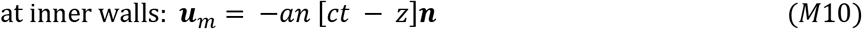

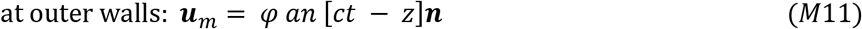

In equations M10 and M11, ‘*an*’ is a periodic function with a time period of 1/f, where f is the heartrate frequency. The waveform of ‘*an*’ is interpolated from the pulsation waveform reported by Mestre et al^7^ (Fig 3b, 4b). The value of *φ* (SAS displacement parameter) is 0 for the simulations presented in Fig 3 and Fig S2. For the simulations presented in Fig 4, the value of *φ* is 0.368.

For the simulations shown in Fig 3, no traction was applied at the axial ends of the PVS (eq. M11–M12). This change was useful to understand the magnitude of pressure changes in the PVS (Fig 3e).

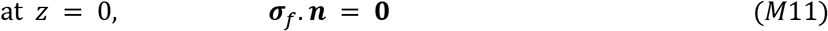

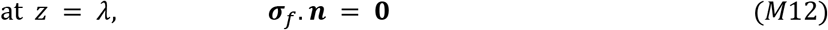

For the simulations shown in Fig 4, no traction is applied at the distal end of the PVS (z = L_a_, where L_a_ is the length of the MCA). This is similar to equation M12. At the proximal, a pressure like traction is applied (eq. M13). The parameter *p*_1_ in equation M11 is the pressure difference across the length of the PVS, shown on the x-axis of Fig 4g. On the peripheral walls of the PVS, the fluid velocity is equal to the wall velocity (similar to eq. M5).

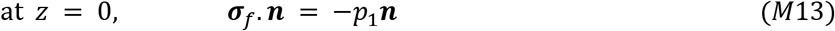

The Reynolds number for all the simulations is calculated using the formula for flow in a pipe (eq. M14). In equation M12, *D_h_* is the hydraulic diameter, which is calculated using the area A and the perimeter P. Q is the flow rate. The Péclet number is calculated using the diffusion (D) coefficient for Amyloid-beta in water (eq. M15). *v_ave_* is the mean downstream speed of the fluid at the center of the PVS (*r* = (*R*_1_ + *R*_2_)/2).

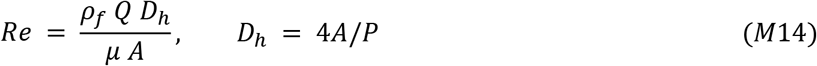

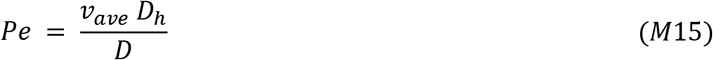

The details about particle tracking in ALE are explained in the appendix. The particle tracking calculations and movies were made using Matlab^®^ code. All the code for Mathematica, Comsol and Matlab are available to download on Github (https://github.com/DrewLab/Peristaltic-pumping-of-CSF.git).

### Anisotropic non-dimensionalization

One of the major concerns when using finite element simulations to study flow in the PVS is the long and narrow geometry of the PVS. For example, the domain used for simulations presented in Fig 2 has a length of one wavelength of the peristaltic wave (116.7 mm or 116,667 μm), which is nearly 3000 times the width of the PVS (40 μm). Simulating the geometry with these dimensions could cause a large number of elements, making it incredibly expensively to solve or create elements with very bad aspect ratios. To deal with this problem, we non-dimensionalized the equations with different scaling factors in the x, y (or r for axisymmetric simulations), and z directions. All the equations from the mesh coordinates (***X**_m_*) are rewritten in these non-dimensional coordinates (***X**_c_*) (eq. M14). In equation M14, the coordinates are written in the conventional order, i.e, (*x, y, z*) for Cartesian and (*r, θ, z*) for cylindrical coordinates.

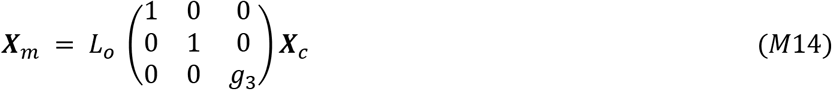

The characteristic length, *L_o_*, was chosen to be equal to the arterial wall radius (R_1_). The scaling factor, *g*_3_, was chosen so that the axial (z) length of the domain in the non-dimensionalized coordinates is 10. This resulted in a scaling factor (*g*_3_) value of ~400 in all the simulations. To verify the validity of this choice of parameter, we plotted the z and r components of the velocity gradient in the mesh coordinates and non-dimensional coordinates (Fig S3). In the mesh coordinates, the velocity gradients (for the radial and the axial component) were nearly three orders of magnitude higher in the radial direction compared to the axial direction. Our choice of scaling factor results in velocity gradients of similar magnitude, which means that for meshes of aspect ratio ~1 in the non-dimensionalized coordinates, the approximation and interpolation errors are rather contained (for low Reynold’s number flows). A similar line of reasoning is used to minimize approximation and interpolation errors for anisotropic adaptive meshing for flow simulations^75–77^.

### Model parameters

All the parameters were taken to match the values observed *in vivo* in mice. The dimensions of the cross-section of the PVS and the pulsation waveform of the arteries were taken from Mestre et al.^7^, to emulate their experimental results. All the parameters used in the model are listed in table 1.

**Table 1.**
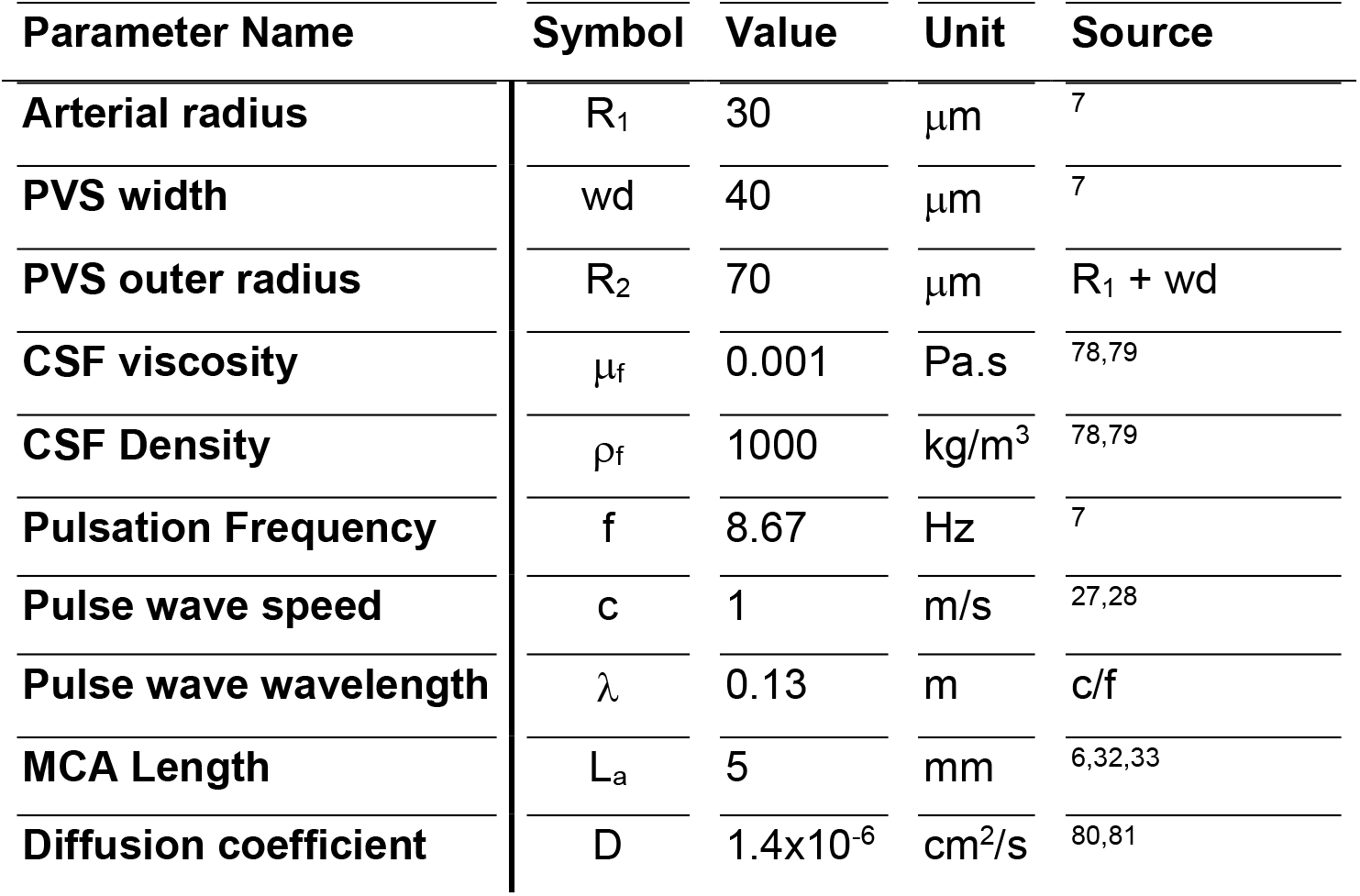
Parameters used in simulations

| Parameter Name | Symbol | Value | Unit | Source |
| --- | --- | --- | --- | --- |
| Arterial radius | $R_1$ | 30 | $\mu\text{m}$ | <sup>7</sup> |
| PVS width | wd | 40 | $\mu\text{m}$ | <sup>7</sup> |
| PVS outer radius | $R_2$ | 70 | $\mu\text{m}$ | $R_1 + \text{wd}$ |
| CSF viscosity | $\mu_f$ | 0.001 | Pa.s | <sup>78,79</sup> |
| CSF Density | $\rho_f$ | 1000 | $\text{kg/m}^3$ | <sup>78,79</sup> |
| Pulsation Frequency | f | 8.67 | Hz | <sup>7</sup> |
| Pulse wave speed | c | 1 | m/s | <sup>27,28</sup> |
| Pulse wave wavelength | $\lambda$ | 0.13 | m | $c/f$ |
| MCA Length | $L_a$ | 5 | mm | <sup>6,32,33</sup> |
| Diffusion coefficient | D | $1.4 \times 10^{-6}$ | $\text{cm}^2/\text{s}$ | <sup>80,81</sup> |

## Acknowledgements

This work was supported by NSF Grant CBET 1705854.

