## Supplementary Figures for "Arterial pulsations drive oscillatory flow of CSF but not directional pumping"

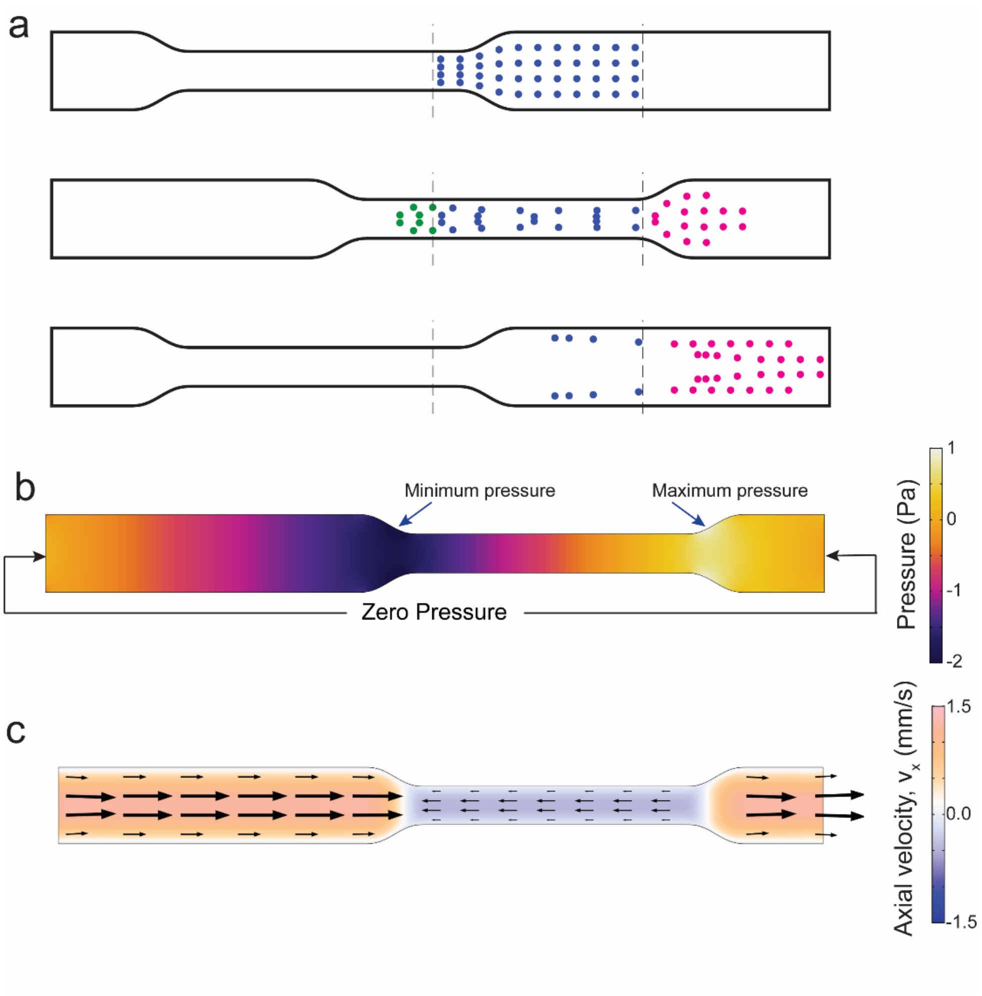


**Figure S1**: Mechanism of fluid motion in a tube driven periodic peristaltic movement of walls.

**a**. The movement of fluid (the dots represent fluid particles) driven by peristaltic compression and relaxation of the walls of a 2D tube. The fluid movement in this figure is calculated using the Navier-Stokes equation with zero traction (pressure) at the two ends of the tube. Fluid in the compressed part of the tube has retrograde flow(middle) but over a full cycle of the peristaltic wave (bottom), the fluid throughout the tube has a net anterograde flow.

**b**. The colors show the pressure in the tube. The cross section of the tube being compressed experiences the maximum pressure. This drives anterograde flow to the right of the cross section and retrograde flow to the left. The anterograde flow is higher than the retrograde flow because it faces a higher flow resistance. This part is same as the mechanism explained in Fig 1a-b. Additionally here, the cross section being relaxed has the minimum pressure and this draws fluid through the left end of the tube. The velocity profile in **c** shows the flow driven by this pressure profile. As the peristaltic wave passes through the tube, the fluid experiences both retrograde flow and anterograde flow with a net flow in the direction of the peristaltic wave.

**c**. The colors show the axial component of fluid velocity ($v_x$). Orange indicates anterograde flow and blue indicates retrograde flow. The arrows are provided to show the direction of flow and make the interpretation of flow easier.

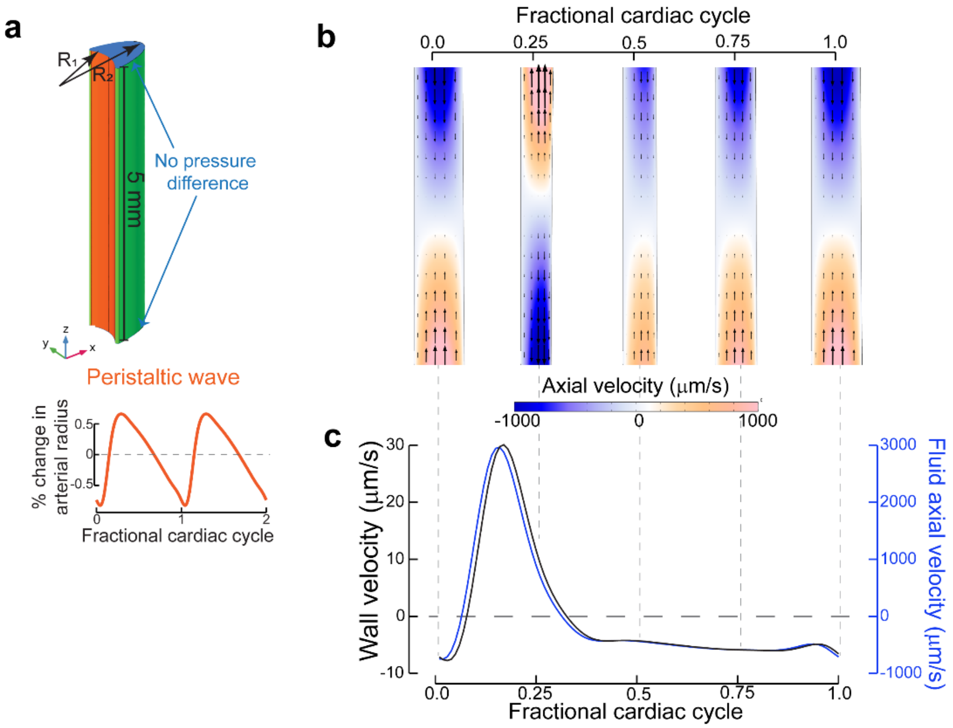


**Figure S2**: Simulations with the correct length of the PVS explain the phase relation between the wall velocity and the fluid velocity

**a**. The dimensions and boundary conditions used in 3D fluid dynamics simulations. The wall movement on the arterial side (orange) is given by a travelling wave with a realistic waveform observed in-vivo in mice[1] (inset). The wall on the brain tissue side is fixed (green). The parameters ($R_1$, $R_2$ are given in Table 1). The length of the PVS is changed from one wavelength of the peristaltic wave (Fig 2) to 5mm to reflect the length of the MCA[2–4] in mice.

**b**. The colors show axial velocity profile at the mid-section of the PVS (XZ plane at y=0) throughout the cardiac cycle. Arrows are provided to make the interpretation of flow easier. Here, the length of the wall is ~20 times smaller than the wavelength of the peristaltic wave, the wall moves in and out almost simultaneously. This means that the cross-sectional area of the tube (and consequently the flow resistance) remains constant throughout the length of the tube resulting in a flow profile very different from the ones expected during peristalsis (Fig 3d). The deformations are increased by a factor of 50 in post-processing to clearly show the arterial wall movement.

**c**. The plots show the arterial wall velocity and the centerline velocity of the fluid taken at the same axial (z = 4mm) location. The peak fluid velocity is in phase with the wall velocity, similar to in-vivo observations[1,2]. The fluid velocities magnitude is 2 orders of magnitude higher than the experimentally observed values. This discrepancy is most likely due to the assumption that the outer walls of the PVS are fixed (see Fig 4e).

The dotted lines between b and c show the shape of the PVS corresponding to the wall and fluid velocity.

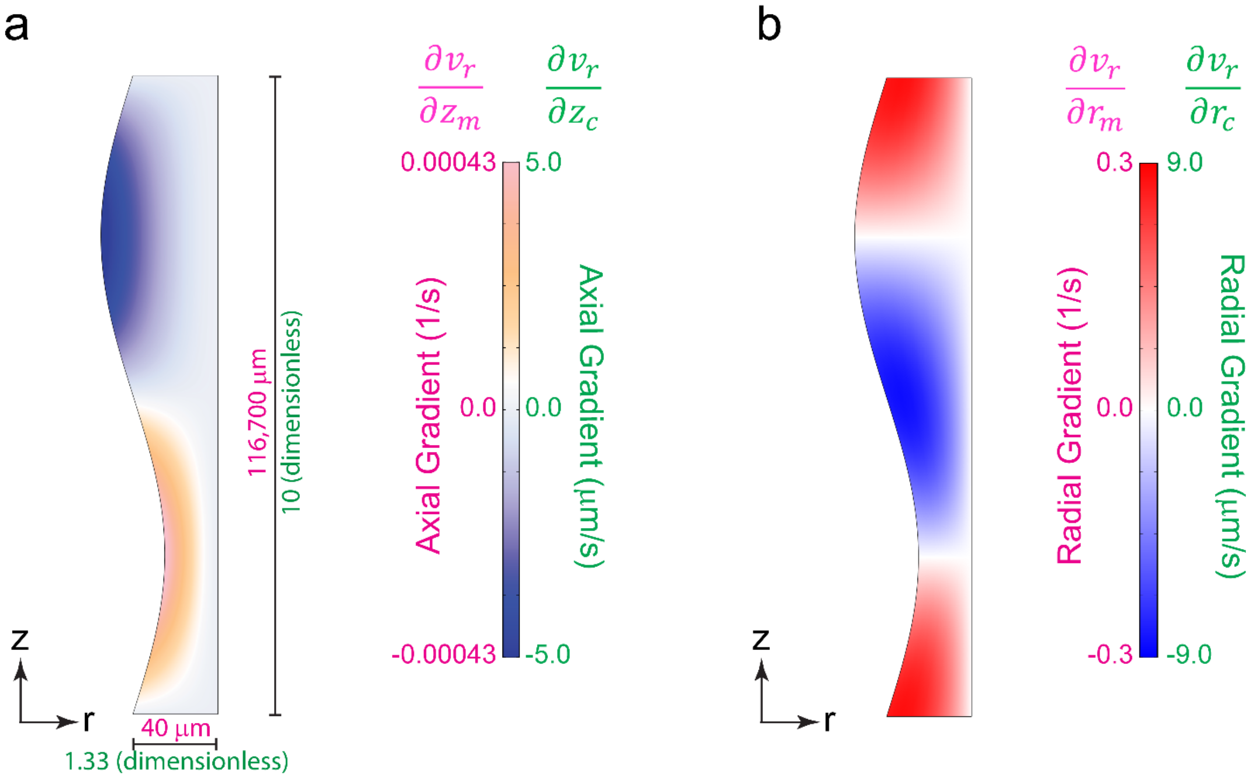


**Figure S3**: Anisotropic non-dimensionalization ensures that the approximation and interpolation errors are minimized

The domain (in the mesh coordinates) has a length to width ratio of ~3000 (**a**). This creates a large difference in the velocity gradients. The component of velocity gradient in the axial direction (**a.** magenta) is 3 orders of magnitude smaller than the component in the radial direction (**b.** magenta). By scaling the geometry anisotropically in the non-dimensional domain, the components of velocity gradient in the radial and axial directions are brought to similar levels (green). By creating meshes of aspect ratio ~1 in the non-dimensionalized domain, we minimize the approximation and interpolation errors.

The results presented in this figure are at the beginning of the cardiac cycle, with a sinusoidal peristaltic wave, whose peak-to-peak amplitude is 1% of the arterial radius.
