## Appendix for "Arterial pulsations drive oscillatory flow of CSF but not directional pumping"

#### A Kinematics in ALE

We start with two configurations of the fluid, the deformed configuration and the reference configuration ($B_t$ and $B_p$ respectively). The deformed configuration ($B_t$) is the image of the reference configuration($B_p$) under the map $\boldsymbol{\chi}_p(\boldsymbol{X}_p, t)$. The points in the reference and deformed configurations are denoted by $\boldsymbol{X}_p, \boldsymbol{x}$ respectively. The displacement of material particles is given by.

$$\boldsymbol{u}(\boldsymbol{X}_p, t) := \boldsymbol{x}(\boldsymbol{X}, t) - \boldsymbol{X}_p \tag{A.1}$$

One can also use the Eulerian description of the displacement

$$\boldsymbol{u}(\boldsymbol{x}, t) := \boldsymbol{x} - \boldsymbol{X}_p(\boldsymbol{x}, t) \tag{A.2}$$

Where $\boldsymbol{X}_p(x, t)$ is the image of $\boldsymbol{x}$ under the inverse map $\boldsymbol{\chi}_p^{-1}(\boldsymbol{x}, t)$.

The velocity $\boldsymbol{v}$ is the time derivative of the displacement. The following relation holds for the Eulerian description of displacement

$$\boldsymbol{v}(\boldsymbol{x}, t) := D_t\boldsymbol{u} \Rightarrow \boldsymbol{v} = \frac{\partial \boldsymbol{u}}{\partial t} + (\boldsymbol{\nabla}_x \boldsymbol{u})\boldsymbol{v} \tag{A.3}$$

where $\boldsymbol{\nabla}_x$ is the gradient with respect to $\boldsymbol{x}$

For calculations in ALE, we define the mesh configuration, where the points are denoted by $\boldsymbol{X}_m$. The deformed configuration $B_t$ is related to the mesh configuration $B_m$ by the map $\chi_m$. The displacement of the mesh domain (mesh motion for finite element calculations) is given by

$$\boldsymbol{x}(\boldsymbol{X}_m, t) - \boldsymbol{X}_m =: \boldsymbol{u}_m(\boldsymbol{X}_m, t) \tag{A.4}$$

The deformation gradient and the Jacobian determinant are defined as

$$\boldsymbol{F}_m(\boldsymbol{X}_m, t) := \frac{\partial \boldsymbol{x}(\boldsymbol{X}_m, t)}{\partial \boldsymbol{X}_m} = \boldsymbol{I} + \boldsymbol{\nabla}_{X_m}\boldsymbol{u}_m \tag{A.5}$$

$$J_m(\boldsymbol{X}_m, t) := \det \boldsymbol{F}_m(\boldsymbol{X}_m, t) \tag{A.6}$$

where, $\boldsymbol{\nabla}_{X_m}$ is the gradient with respect to $\boldsymbol{X}_m$ over $B_m$.

The primary unknown fields,for example, the stress tensor($\boldsymbol{\sigma}$) fluid velocity ($\boldsymbol{v}$) and pressure ($p$) in the mesh coordinates $\boldsymbol{X}_m$ are given by:

$$\hat{p}(\boldsymbol{X}_m, t) = p(\boldsymbol{x}, t) = p(\boldsymbol{x}(\boldsymbol{X}_m, t), t) \tag{A.7}$$

$$\hat{\boldsymbol{v}}(\boldsymbol{X}_m, t) = \boldsymbol{v}(\boldsymbol{x}, t) = \boldsymbol{v}(\boldsymbol{x}(\boldsymbol{X}_m, t), t) \tag{A.8}$$

$$\hat{\boldsymbol{\sigma}}(\boldsymbol{X}_m, t) = \boldsymbol{\sigma}(\boldsymbol{x}, t) = \boldsymbol{\sigma}(\boldsymbol{x}(\boldsymbol{X}_m, t), t) \tag{A.9}$$

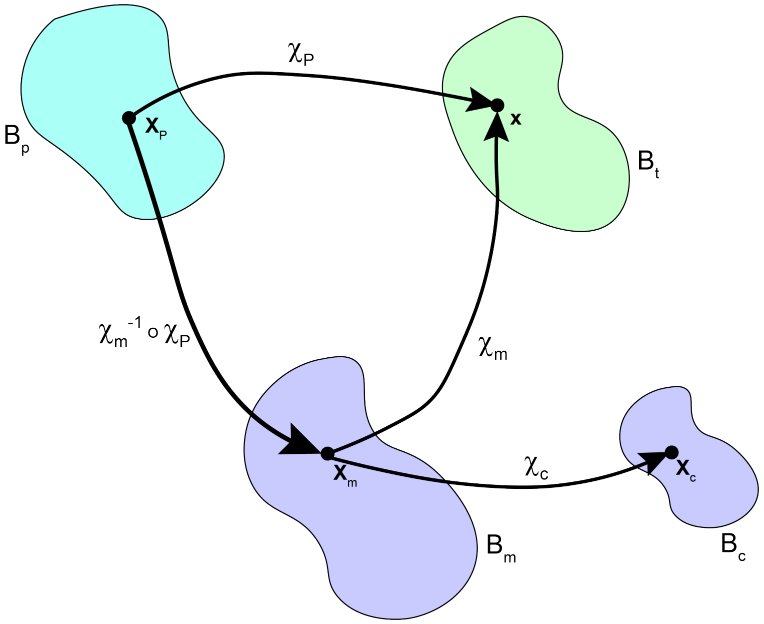


Figure A.1: The current (deformed) configuration, $B_t$, and it's relation to the reference configuration ($B_p$) and the mesh configuration ($B_m$). The computational configuration ($B_c$) is an anisotropically non-dimensionalized transformation of the mesh configuration.

The material time derivative of $\boldsymbol{x}(\boldsymbol{X}_m, t)$ is the material particle velocity in mesh Coordinates, which should be the same as the material particle velocity defined in eq. A.3

$$D_t\boldsymbol{x}(\boldsymbol{X}_m, t) =: \hat{\boldsymbol{v}}(\boldsymbol{X}_m, t) = \boldsymbol{v}(\boldsymbol{x}(\boldsymbol{X}_m, t), t) \tag{A.10}$$

The deformation gradient defined in eq. A.5 and the Jacobian determinant defined in eq. A.6 are used for transforming the spatial derivatives to the mesh domain.

For a vector $\boldsymbol{v}$:

$$\boldsymbol{\nabla}_x\boldsymbol{v} = \boldsymbol{\nabla}_{X_m}\hat{\boldsymbol{v}}\boldsymbol{F}_m^{-1} \tag{A.11}$$

$$\boldsymbol{\nabla}_x.\boldsymbol{v} = Tr[\boldsymbol{\nabla}_x\boldsymbol{v}] = Tr[\boldsymbol{\nabla}_{X_m}\hat{\boldsymbol{v}}\boldsymbol{F}_m^{-1}] \tag{A.12}$$

where, Tr is the trace operator on tensors.

For a scalar $p$:

$$\boldsymbol{\nabla}_x p = \boldsymbol{F}_m^{-T}\boldsymbol{\nabla}_{X_m}\hat{p} \tag{A.13}$$

For a tensor $\boldsymbol{\sigma}$

$$\boldsymbol{\nabla}_x.\boldsymbol{\sigma} = \frac{1}{J_m}\boldsymbol{\nabla}_{X_m}.\left(J_m\hat{\boldsymbol{\sigma}}\boldsymbol{F}_m^{-T}\right) \tag{A.14}$$

The material time derivative of velocity(the acceleration) is calculated as

$$\hat{\boldsymbol{a}}(\boldsymbol{X}_m, t) := D_t\hat{\boldsymbol{v}}(\boldsymbol{X}_m, t) = \frac{\partial\hat{\boldsymbol{v}}}{\partial t} + (\boldsymbol{\nabla}_{X_m}\hat{\boldsymbol{v}})\boldsymbol{F}_m^{-1}\left(\hat{\boldsymbol{v}} - \frac{\partial\boldsymbol{u}_m}{\partial t}\right) \tag{A.15}$$

A detailed derivation of eq. A.15 was done by Donea *et al.* [5].

Integrals over the deformed domain can be transformed to the mesh domain as follows

$$\int_{B_t} f(\boldsymbol{x})\,\mathrm{d}\nu = \int_{B_m} J_m \hat{f}(\boldsymbol{X}_m)\,\mathrm{d}\nu_m \tag{A.16}$$

where $\nu$ and $\nu_m$ are infinitesimal volumes is $B_t$ and $B_m$ respectively.

Integrals over the boundaries are transformed as follows

$$\int_{\partial B_t} \boldsymbol{g}(\boldsymbol{x}).\boldsymbol{n}\,\mathrm{d}a = \int_{\partial B_m} J_m \hat{\boldsymbol{g}}(\boldsymbol{X}_m).\boldsymbol{F}_m^{-T}\boldsymbol{n}_m\,\mathrm{d}a_m \tag{A.17}$$

where $a$ and $a_m$ are infinitesimal areas in $\partial B_t$ and $\partial B_m$ respectively and $\boldsymbol{n}_m$ is a unit normal in the mesh domain.

### B Anisotropic non-dimensionalization

We transform the mesh domain into the non-dimensionalized computational domain with coordinates $\hat{\boldsymbol{X}}_c$. The transformation is defined by the characteristic length $L_o$ and scaling tensor $\boldsymbol{G}$. For our problem, the characteristic length and the scaling tensor are constants.

$$\boldsymbol{X}_m = L_o \boldsymbol{G} \boldsymbol{X}_c \tag{B.1}$$

Any field in $X_m$ can be transformed into a field in $X_c$ using the relation in B.1

$$f(\boldsymbol{X}_m) = f(L_o \boldsymbol{G} \boldsymbol{X}_c) \tag{B.2}$$

The spatial derivatives are transformed as follows.

For a vector $\boldsymbol{v}$:

$$\boldsymbol{\nabla}_{X_m}\boldsymbol{v} = \frac{1}{L_o}\boldsymbol{\nabla}_{X_c}\boldsymbol{v}\boldsymbol{G}^{-1} \tag{B.3}$$

$$\boldsymbol{\nabla}_{X_m}.\boldsymbol{v} = Tr[\boldsymbol{\nabla}_{X_m}\boldsymbol{v}] = \frac{1}{L_o}Tr[\boldsymbol{\nabla}_{X_c}\boldsymbol{v}\boldsymbol{G}^{-1}] \tag{B.4}$$

For a scalar $p$:

$$\boldsymbol{\nabla}_{X_m}p = \frac{1}{L_o}\boldsymbol{G}^{-T}\boldsymbol{\nabla}_{X_c}p \tag{B.5}$$

For a tensor $\boldsymbol{\sigma}$:

$$\boldsymbol{\nabla}_{X_m}.\boldsymbol{\sigma} = \frac{1}{L_o}\boldsymbol{\nabla}_{X_c}.\Big(\boldsymbol{\sigma}\boldsymbol{G}^{-T}\Big) \tag{B.6}$$

The time derivatives are unaltered because there is no relative motion between $X_m$ and $X_c$.

The volume and surface integrals are transformed as follows.

$$\int_{B_m} f(\boldsymbol{X}_m)\,\mathrm{d}\nu_m = L_o^3 \det[\boldsymbol{G}] \int_{B_c} f(L_o\boldsymbol{G}\boldsymbol{X}_c)\,\mathrm{d}\nu_c \tag{B.7}$$

Integrals over the boundaries are transformed as:

$$\int_{\partial B_m} \boldsymbol{g}(\boldsymbol{X}_m).\boldsymbol{n}_m\,\mathrm{d}a_m = L_o^2 \det[\boldsymbol{G}] \int_{\partial B_c} \boldsymbol{g}(L_o\boldsymbol{G}\boldsymbol{X}_c).\boldsymbol{G}^{-T}\boldsymbol{n}_c\,\mathrm{d}a_c \tag{B.8}$$

#### C Initial conditions for problems

In all the intial-boundary value problems used in the study, the initial values and initial time-derivatives of all variables are set to zero because the initial conditions of the system are unknown. The parameter values used in the boundary conditions, are ramped up from zero to the specified values using step functions available in Comsol. These functions have continuous first and second derivatives with respect to time. In this document we will use $step(t)$ in the equations to indicate wherever these functions are used. The results for these simulations are shown after 4 cycles of the periodic wall motion, where the variations in the velocity and pressure values from cycle to cycle are less than $10^{-4}\%$ of peak value.

#### D Navier-Stokes in ALE coordinates

We solve for the fluid velocity $\boldsymbol{v}_f$ and pressure $p_f$ in a deforming domain $B_t$, representing the PVS. The time-dependent deformation of the domain is represented by $\boldsymbol{u}_m$ (same as eq. A.4).We present the strong form equations here in the mesh coordinates $\boldsymbol{X}_m$.The equations can be verified with previously published literature[3, 2].The appropriate transformations for change of coordinates are applied (eq. A.11 - eq. A.17) and the $X_m$ subscript on the spatial derivatives (gradient, divergence and laplacian) is dropped.

We use a linear elliptical model for the mesh motion [1, 4].The governing equation for the mesh displacement, $\boldsymbol{u}_m$ on $B_m$ is given by

$$\nabla^2 \boldsymbol{u}_m = 0 \tag{D.1}$$

The equations in the mesh coordinates are written for the velocity and pressure in the mesh coordinates($\hat{\boldsymbol{v}}_f$ and $\hat{p}_f$). The governing equation for the fluid in the PVS(Navier-Stokes) are:

$$\frac{\partial \hat{\boldsymbol{v}}_f}{\partial t} + \boldsymbol{\nabla} \hat{\boldsymbol{v}}_f \boldsymbol{F}_m^{-1} \left( \hat{\boldsymbol{v}}_f - \frac{\partial \boldsymbol{u}_m}{\partial t} \right) - \frac{1}{J_m \rho_f} \boldsymbol{\nabla}.(J_m \hat{\boldsymbol{\sigma}}_f \boldsymbol{F}_m^{-T}) = \boldsymbol{0} \tag{D.2}$$

$$Tr[\boldsymbol{\nabla} \hat{\boldsymbol{v}}_f \boldsymbol{F}_m^{-1}] = 0 \tag{D.3}$$

$$\hat{\boldsymbol{\sigma}}_f = -\hat{p}_f \boldsymbol{I} + \mu_f \left( \boldsymbol{\nabla} \hat{\boldsymbol{v}}_f \boldsymbol{F}_m^{-1} + \left( \boldsymbol{\nabla} \hat{\boldsymbol{v}}_f \boldsymbol{F}_m^{-1} \right)^T \right) \tag{D.4}$$

The Dirichlet boundary conditions specified in equations M5-M8,M10 and M11 (see methods) are applied as-is. The zero traction boundary conditions in the form of equation M12 and M13 are applied as equation D.5 and the pressure-like traction is applied as equation D.6

$$J_m \hat{\boldsymbol{\sigma}}_f \boldsymbol{F}_m^{-T} \boldsymbol{n}_m = \boldsymbol{0} \tag{D.5}$$

$$J_m \hat{\boldsymbol{\sigma}}_f \boldsymbol{F}_m^{-T} \boldsymbol{n}_m = -J_m p_1 \boldsymbol{F}_m^{-T} \boldsymbol{n}_m \tag{D.6}$$

#### E Weak form equations in non-dimensional coordinates

The finite element calculations are performed in the anisotropically non-dimensionalized computational coordinates. All the equations in this section are expressed in the computational coordinates($\boldsymbol{X}_c$).The appropriate transformations listed in equations B.2 - B.8 are applied and the subscript $\boldsymbol{X}_c$ is dropped.

The boundary of the computational domain is divided into three subsets, $\Gamma^D$, $\Gamma^P$ and $\Gamma^N$, representing the surfaces with Dirichlet, Periodic and Neumann Boundary conditions respectively. The periodic boundary, $\Gamma^P$, is further divided into $\Gamma^{P1}$ and $\Gamma^{P2}$, representing the source and destination for the periodic boundary conditions. We define the following functional spaces in the computational domain ($B_c$) for velocity, pressure and mesh displacement respectively.

$$\mathcal{V}^{\boldsymbol{v}_f} := \left\{ \boldsymbol{v}_f \in L^2(B_c)^d \mid \boldsymbol{\nabla} \boldsymbol{v}_f \in L^2(B_c)^{d\times d}, \boldsymbol{v}_f = \bar{\boldsymbol{v}}_f \ on\ \Gamma^D, \boldsymbol{v}_f|_{\Gamma^{P1}} = \boldsymbol{v}_f|_{\Gamma^{P2}} \right\} \tag{E.1}$$

$$\mathcal{V}^{p_f} := \left\{ p_f \in L^2(B_c)^d \mid \boldsymbol{\nabla} p_f \in L^2(B_c)^{d\times d}, p_f|_{\Gamma^{P1}} = p_f|_{\Gamma^{P2}} \right\} \tag{E.2}$$

$$\mathcal{V}^{\boldsymbol{u}_m} := \left\{ \boldsymbol{u}_m \in L^2(B_c)^d \mid \boldsymbol{\nabla} \boldsymbol{u}_m \in L^2(B_c)^{d\times d}, \boldsymbol{u}_m = \bar{\boldsymbol{u}}_m \ on\ \Gamma^D, \boldsymbol{u}_m|_{\Gamma^{P1}} = \boldsymbol{u}_m|_{\Gamma^{P2}} \right\} \tag{E.3}$$

Additionally, we define the spaces

$$\mathcal{V}_0^{\boldsymbol{v}_f} := \left\{\boldsymbol{v}_f \in L^2(B_c)^d \mid \boldsymbol{\nabla}\boldsymbol{v}_f \in L^2(B_c)^{d\times d}, \boldsymbol{v}_f = \boldsymbol{0} \ on \ \Gamma^D \cup \Gamma^{P2}\right\} \tag{E.4}$$

$$\mathcal{V}_0^{p_f} := \left\{p_f \in L^2(B_c)^d \mid \boldsymbol{\nabla} p_f \in L^2(B_c)^{d\times d}, p_f = 0 \ on \ \Gamma^{P2}\right\} \tag{E.5}$$

$$\mathcal{V}_0^{\boldsymbol{u}_m} := \left\{\boldsymbol{u}_m \in L^2(B_c)^d \mid \boldsymbol{\nabla}\boldsymbol{u}_m \in L^2(B_c)^{d\times d}, \boldsymbol{u}_m = \boldsymbol{0} \ on \ \Gamma^D \cup \Gamma^{P2}\right\} \tag{E.6}$$

In equations E.1 to E.6, d is the number of dimensions of the problem. The abstract weak formulation of the problem is as follows:
Find $\hat{\boldsymbol{v}}_f \in \mathcal{V}^{\boldsymbol{v}_f}$, $\hat{p}_f \in \mathcal{V}^{p_f}$ and $\boldsymbol{u}_m \in \mathcal{V}^{\boldsymbol{u}_m}$, such that $\forall$ $\tilde{\boldsymbol{v}}_f \in \mathcal{V}_0^{\boldsymbol{v}_f}$, $\tilde{p}_f \in \mathcal{V}_0^{p_f}$ and $\tilde{\boldsymbol{u}}_m \in \mathcal{V}_0^{\boldsymbol{u}_m}$

$$\int_{B_c} J_m \tilde{\boldsymbol{v}}_f . \left(\frac{\partial \hat{\boldsymbol{v}}_f}{\partial t} + \frac{1}{L_o}\boldsymbol{\nabla}\hat{\boldsymbol{v}}_f \boldsymbol{G}^{-1}\boldsymbol{F}_m^{-1}\left(\hat{\boldsymbol{v}}_f - \frac{\partial \boldsymbol{u}_m}{\partial t}\right)\right) + \int_{B_c} \frac{J_m}{L_o \rho_f}\boldsymbol{\nabla}\tilde{\boldsymbol{v}}_f : \boldsymbol{\sigma}_f \boldsymbol{F}_m^{-T}\boldsymbol{G}^{-T} = 0 \tag{E.7}$$

$$\int_{B_c} J_m \tilde{p}_f Tr\left[\boldsymbol{\nabla}\hat{\boldsymbol{v}}_f \boldsymbol{G}^{-1}\boldsymbol{F}_m^{-1}\right] = 0 \tag{E.8}$$

$$\int_{B_c} \boldsymbol{\nabla}\tilde{\boldsymbol{u}}_m \boldsymbol{G}^{-1} : \boldsymbol{\nabla}\boldsymbol{u}_m \boldsymbol{G}^{-1} = 0 \tag{E.9}$$

$$where, \ \hat{\boldsymbol{\sigma}}_f = -\hat{p}_f \boldsymbol{I} + \frac{\mu_f}{L_o}\left(\boldsymbol{\nabla}\hat{\boldsymbol{v}}_f \boldsymbol{G}^{-1}\boldsymbol{F}_m^{-1} + \left(\boldsymbol{\nabla}\hat{\boldsymbol{v}}_f \boldsymbol{G}^{-1}\boldsymbol{F}_m^{-1}\right)^T\right) \tag{E.10}$$

For cases where there are surfaces with a non-zero traction boundary conditions($\Gamma^{D1}$), an additional term $TBC_f$(eq. E.11) is added to left side of equation E.7.

$$TBC_f = \int_{\Gamma^{D1}} p_1 \frac{J_m}{\rho_f L_o}\tilde{\boldsymbol{v}}_f . F_m^{-T}\boldsymbol{G}^{-T}\boldsymbol{n}_c \tag{E.11}$$

### F Particle tracking in ALE

To find the fluid particle trajectories in the deforming domain, $B_t$, we first solve for the fluid particle coordinates in the mesh domain ($\boldsymbol{X}_m$) as a function of time. The material time derivative of $\boldsymbol{X}_m$ is the particle velocity observed from the mesh domain $B_m$ is defined as:

$$D_t \boldsymbol{X}_m =: \dot{\boldsymbol{X}}_m \tag{F.1}$$

We need to calculate the material time derivative of the mesh displacement. We use a method similar to that in eq. A.3, where we use the map $\chi_m^{-1} o \chi_p$ instead of the map $\chi_p$ in eq.A.3.

$$D_t \boldsymbol{u}_m(\boldsymbol{X}_m, t) = \frac{\partial \boldsymbol{u}_m}{\partial t} + (\boldsymbol{\nabla}_{X_m}\boldsymbol{u}_m)\dot{\boldsymbol{X}}_m \tag{F.2}$$

The finite element computations in the computational coordinates give us the fields $\hat{\boldsymbol{v}}(\boldsymbol{X}_c, t)$ and $\boldsymbol{u}_m(\boldsymbol{X}_c, t)$. These fields can be transformed into mesh coordinates using the inverse of the transformation in eq. B.2. We calculate $\dot{\boldsymbol{X}}_m$ as a function of these known quantities using equations A.10, F.1 and F.2 in the material time derivative of eq. A.4.

$$\hat{\boldsymbol{v}} - \dot{\boldsymbol{X}}_m = \frac{\partial \boldsymbol{u}_m}{\partial t} + (\boldsymbol{\nabla}_{X_m}\boldsymbol{u}_m)\dot{\boldsymbol{X}}_m \tag{F.3}$$

$$(\boldsymbol{I} + \boldsymbol{\nabla}_{X_m}\boldsymbol{u}_m)\dot{\boldsymbol{X}}_m = \hat{\boldsymbol{v}} - \frac{\partial \boldsymbol{u}_m}{\partial t} \tag{F.4}$$

Using the definition of $\boldsymbol{F}_m$ from eq. A.5 we obtain the particle velocity in the mesh domain

$$\dot{\boldsymbol{X}}_m = \boldsymbol{F}_m^{-1}\left(\hat{\boldsymbol{v}} - \frac{\partial \boldsymbol{u}_m}{\partial t}\right) \tag{F.5}$$

We use the following initial value problem to calculate fluid particle trajectories the deforming domain. Here, it is assumed that the material particle velocity $\hat{\boldsymbol{v}}(\boldsymbol{X}_m, t)$ and the mesh displacement $\boldsymbol{u}_m(\boldsymbol{X}_m, t)$ (and subsequently $\boldsymbol{F}_m$ and $\frac{\partial \boldsymbol{u}_m}{\partial t}$) are known fields in the mesh domain. We first find the particle trajectory in the mesh coordinates and add the mesh displacement to find the particle trajectory in the deformed coordinates.

$$\textit{find } \boldsymbol{x}(\boldsymbol{X}_m, t) \textit{ such that} \begin{cases} \boldsymbol{x}(\boldsymbol{X}_m, t) & = \boldsymbol{X}_m + \boldsymbol{u}_m(\boldsymbol{X}_m, t) \\ \dot{\boldsymbol{X}}_m & = \boldsymbol{F}_m^{-1}\left(\hat{\boldsymbol{v}}(\boldsymbol{X}_m, t) - \frac{\partial \boldsymbol{u}_m(\boldsymbol{X}_m, t)}{\partial t}\right) \end{cases} \tag{F.6}$$

With the initial condition: $\boldsymbol{x}(\boldsymbol{X}_m, 0) = \boldsymbol{X}_m + \boldsymbol{u}_m(\boldsymbol{X}_m, 0)$

We solve this problem using a forward Euler integration scheme and calculate the fluid particle trajectories. When the position in the mesh domain at time t is known, the time integration scheme allows us to find the position at time t+dt.

$$\boldsymbol{X}_m(t + dt) = \boldsymbol{X}_m(t) + dt\boldsymbol{F}_m(\boldsymbol{X}_m(t), t)^{-1}\left(\hat{\boldsymbol{v}}(\boldsymbol{X}_m(t), t) - \frac{\partial \boldsymbol{u}_m(\boldsymbol{X}_m(t), t)}{\partial t}\right) \tag{F.7}$$

The fluid particles can only be tracked till the time they exit the computational domain since the velocity field outside is unknown.
